# A mobile intron facilitates interference competition between co-infecting viruses

**DOI:** 10.1101/2023.09.30.560319

**Authors:** Erica A. Birkholz, Chase J. Morgan, Thomas G. Laughlin, Rebecca K. Lau, Amy Prichard, Sahana Rangarajan, Gabrielle N. Meza, Jina Lee, Emily G. Armbruster, Sergey Suslov, Kit Pogliano, Justin R. Meyer, Elizabeth Villa, Kevin D. Corbett, Joe Pogliano

## Abstract

Mobile introns containing homing endonucleases are widespread in nature and have long been assumed to be selfish elements that provide no benefit to the host organism. These genetic elements are common in viruses, but whether they confer a selective advantage is unclear. Here we studied a mobile intron in bacteriophage ΦPA3 and found its homing endonuclease gp210 contributes to viral competition by interfering with the virogenesis of co-infecting phage ΦKZ. We show that gp210 targets a specific sequence in its competitor ΦKZ, preventing the assembly of progeny viruses. This work reports the first demonstration of how a mobile intron can be deployed to engage in interference competition and provide a reproductive advantage. Given the ubiquity of introns, this selective advantage likely has widespread evolutionary implications in nature.

## Introduction

Mobile introns are widespread in all kingdoms of life^1–5^. Despite their ubiquity, it remains unclear whether these genetic elements are wholly selfish or whether they may also provide a selective advantage to their host^6–10^. Mobile introns in viruses have been shown to influence the fitness of neighboring genes, in a process termed marker exclusion^7,11–16^. However, there has not been any reported competitive advantage for a virus that is conferred by a mobile intron.

Previously, we characterized the first known intracellular speciation factors that separate the gene pools of co-infecting phages by studying the nucleus-forming jumbo phages ΦKZ and ΦPA3 of *Pseudomonas aeruginosa*^17^. These phages form a proteinaceous phage nucleus^18–20^ that shields their DNA genomes from host defenses^21–23^ and, like the eukaryotic nucleus, strictly separates transcription and DNA replication from translation and metabolic processes. A protein naturally imported into the ΦPA3 nucleus, gp210, is excluded from the ΦKZ nucleus^17^. To test the outcome of artificially importing gp210 into the ΦKZ nucleus, we utilized our discovery that GFPmut1 is imported into the ΦKZ nucleus while other variants of GFP, such as sfGFP, are not imported^17,21^. When gp210 is fused to GFPmut1, it is imported into the ΦKZ nucleus, resulting in a significant inhibition of plaque formation^17^ with a 0.0017% efficiency of plating (EOP) compared to control cells expressing only GFPmut1 (Figure 1A). Given that ΦPA3 and ΦKZ can coinfect the same cell and assemble a hybrid nucleus^17^, we reasoned that ΦPA3 gp210 might interfere with ΦKZ reproduction during natural co-infections thereby providing a selective advantage to ΦPA3. We therefore investigated the molecular basis of this effect.

**Figure 1.**
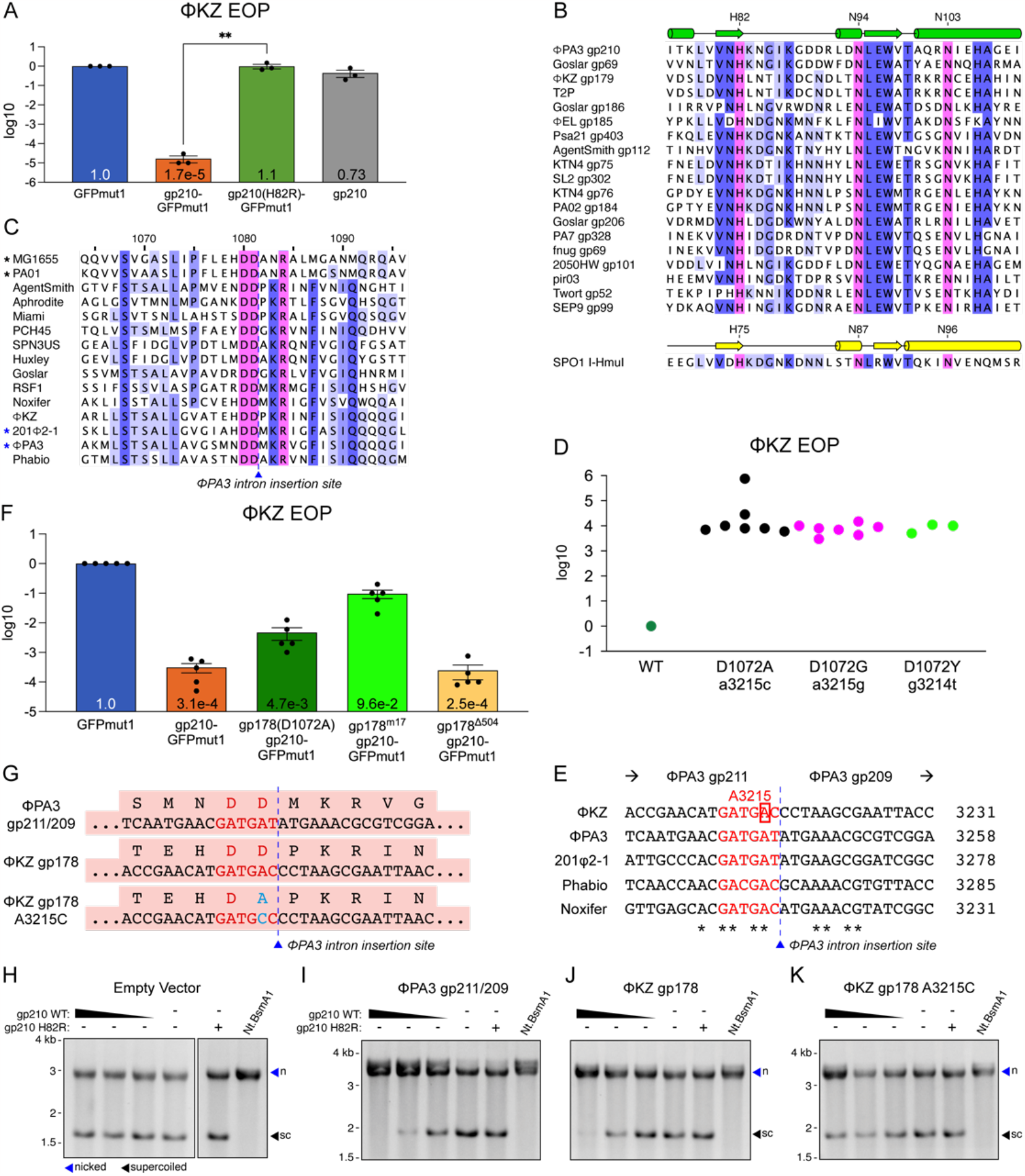
Gp210 of ΦPA3 is an HNH homing endonuclease. (A) Efficiency of plating (EOP) of ΦKZ as measured by spot titer and normalized to the paired titer on GFPmut1. gp210-GFPmut1 causes a 99.9983% decrease in ΦKZ titer (*p=0*.*0011*, n=3) while untagged gp210, not imported into the nucleus, causes an insignificant 27% decrease (*p=0*.*1371*, n=3). H82R mutation (gp210(H82R)-GFPmut1) fully rescues ΦKZ titer compared to gp210-GFPmut1 (*p=0*.*0026*, n=3). Error bars represent SEM, p values calculated by ratio paired t-test. (B) Protein alignment of phage-encoded endonucleases related to gp210 (top is predicted secondary structure from Phyre2), including well-characterized homing endonuclease I-HmuI (bottom with confirmed secondary structure). ClustalO alignment. (C) Protein alignment including the intron(-) version of the ΦPA3 RNAP, after editing from the annotation in Genbank which can be found in Figure S1D. It is aligned with RpoB of MG1655 (*E. coli*) and PAO1 (*P. aeruginosa*), 8 RNAPs encoded by jumbo phages infecting different genera of hosts, and 4 RNAPs encoded by other *Pseudomonas* jumbo phages. ClustalO alignment. The intron insertion site occupied in the ΦPA3 gene is indicated. Residue numeration above is based on the ΦPA3 sequence. (D) Efficiency of plating (EOP) of ΦKZ escape mutants relative to the wildtype ΦKZ on cells expressing gp210-GFPmut1. (E) Nucleotide alignment of intron(-) RpoB homologs in 5 *Pseudomonas* jumbo phages. The point mutation in the ΦKZ vRNAP gp178 at position 3215 (red box) is 2 bases upstream of the site that aligns with the intron insertion site (IIS, black dotted line) in ΦPA3. Red sequences code for the conserved aspartic acid residues. The exons of ΦPA3 are noted above. (F) EOP of ΦKZ on cells expressing GFPmut1 as a control, gp210-GFPmut1, or co-expressing gp210-GFPmut1 and one of three variations on the gp178 ΦKZ vRNAP gene. The nucleotide sequences of these variants are diagrammed in Figure S2F. (G) Sequences surrounding the insertion site of the intron containing gp210. ΦPA3 gp211/209: intron(-) version of the ΦPA3 RNAP subunit. ΦKZ gp178: homologous RNAP subunit. ΦKZ gp178 A3215C: gp178 with single nucleotide mutation. (H-K) Nuclease assay of purified gp210 incubated with plasmid DNA containing only the empty vector (H), the intron(-) PA3 gene (I, gp211/209), KZgp178 gene (J) or KZgp178 gene with nucleotide change A3215C (K). Gp210 concentration from left to right was 5.0, 2.5, 1.25, and 0 μM. Nt.BsaI is a reference digest for nicked plasmid (blue arrows, n). Supercoiled plasmid is indicated by black arrows, sc.

## Results

Gp210 is predicted to be an HNH endonuclease based on sequence alignments and AlphaFold structure predictions (Figures 1B and S1A-D). We mutated gp210’s predicted active site histidine to arginine (H82R), and found that expression of gp210(H82R)-GFPmut1 in host *P. aeruginosa* cells was no longer able to reduce ΦKZ’s viral titer (Figure 1A), despite being successfully imported into the ΦKZ nucleus (Figure S1F). Confirming the likely role of gp210 nuclease activity in ΦKZ suppression, a 1,000-fold higher multiplicity of infection (MOI) of ΦKZ was required to suppress the growth of *P. aeruginosa* cells expressing wild-type gp210-GFPmut1 compared to cells expressing gp210(H82R)-GFPmut1 (Figure S1E). In contrast, ΦPA3 was unaffected by expression of either gp210-GFPmut1 or gp210(H82R)-GFPmut1 (Figures S2A-C).

Gp210 is encoded within a putative group I self-splicing intron. Homing endonucleases, such as SPOI I-HmuI, are often encoded within self-splicing group I introns that have inserted into a highly conserved region of an essential gene, and can target their own insertion site locus as well as a homologous locus of co-infecting phages^16,24–27^. Gp210 interrupts the gene encoding the β subunit of the virion-packaged ΦPA3 RNA polymerase (virion RNA polymerase or vRNAP), and is inserted immediately following two highly conserved aspartic acid residues (Figure 1C), the second of which has been implicated in polymerase fidelity of RpoB in *E. coli* (D675)^28^. To determine whether gp210 is a nuclease that targets this conserved region within the ΦKZ genome, we analyzed ΦKZ escaper mutants that were able to infect *P. aeruginosa* expressing gp210-GFPmut1 (Figures 1D and E). We isolated fourteen spontaneous escaper mutants that had a mutation of the adenosine at position 3215 of the gp178 gene, resulting in a D1072A or D1072G amino acid change in the resulting vRNAP subunit (Figure 1D). A further three escapers mutated the guanine at position 3214, resulting in a gp178 D1072Y mutation. The sequence surrounding these sites is highly conserved among ΦKZ-related phages (Figure 1E). All of these changes resulted in a missense mutation of the highly conserved aspartic acid residue at position 1072 to alanine, glycine, or tyrosine. D1072 of ΦKZ gp178 aligns with the second aspartic acid at the end of ΦPA3 gp211, which borders the gp210 intron insertion site in ΦPA3 (Figures 1C and G).

To confirm that gp210 is a homing endonuclease that targets the intron(-) version of its genomic locus in ΦPA3 and the homologous gene in ΦKZ, we created plasmids containing the uninterrupted sequence of the insertion site in the ΦPA3 RNAP subunit (ΦPA3 gp211/209) or the matching locus in the homologous ΦKZ gene 178 and tested the nucleolytic activity of purified gp210 *in vitro* (Figures 1H-K). While the empty vector was not cleaved by gp210, both the intron-deleted ΦPA3 gp211/209 sequence and the ΦKZ gp178 sequence were cleaved to convert supercoiled plasmid to open circular, similar to the Nt.BsmA1 nicking control enzyme (Figures 1H-J). This activity required the HNH domain, as the H82R mutant protein showed no DNA cleavage. We tested whether ΦPA3 gp210 could cleave a modified ΦKZ gp178 sequence with the A3215C mutation, and found that cleavage was reduced (Figure 1K). This demonstrates that gp210 is capable of cutting the gene encoding ΦKZ gp178, a vRNAP subunit that is packaged into ΦKZ virions^29^ and transcribes early genes^30^.

The above results suggest that gp210 cleaves the ΦKZ genome within the gp178 gene, and that either the loss of this gene product or the lost integrity of the genomic DNA itself is responsible for the ΦPA3 gp210-dependent knockdown of ΦKZ viral titer. To distinguish between these hypotheses, we determined whether expression of gp178 on a plasmid can rescue ΦKZ replication from gp210-GFPmut1. We co-expressed gp210-GFPmut1 with either gp178(D1072A) containing adenosine 3215 mutated to cytosine or gp178 with seventeen silent mutations near the putative gp210 cleavage site to prevent gp210 recognition but conserve the amino acid sequence (gp178^m17^) (Figure S2F). We found that co-expression of gp178(D1072A) resulted in a 15-fold increase in ΦKZ titer compared to only gp210-GFPmut1 and co-expression of gp178^m17^ resulted in a 300-fold increase compared to only gp210-GFPmut1 (Figure 1F). Co-expression of gp178 with a large internal in-frame deletion of 504 bases (gp178^Δ504^) did not rescue ΦKZ, demonstrating that co-expression of gp210-GFPmut1 downstream of another ORF is not responsible for the rescue. Consistent with this region being the recognition sequence for gp210, we could not construct a plasmid for co-expression of gp210-GFPmut1 and the wildtype gp178 gene sequence. Taken together, these results support the hypothesis that gp210 of ΦPA3 is able to hydrolyze ΦKZ DNA within the vRNAP gene gp178. The proximity of the mutation to the native intron insertion site of gp210 also indicates typical homing activity.

Despite a drastic inhibition of ΦKZ titer in cells expressing gp210-GFPmut1, the phage nucleus still formed, enlarged, and was centered with bright DAPI staining (Figure 2A). Normally at 70 mpi, DNA-containing ΦKZ particles assemble into bouquet structures that can be visualized with DAPI staining^31^. However, expression of gp210-GFPmut1 resulted in a lack of stained capsids in the ΦKZ phage bouquets by 70 mpi (Figure 2A yellow arrowheads). This change in the distribution of DAPI staining was measured by line plots of DAPI intensity along a bisecting line (Figure 2C, n=50), supporting the conclusion that capsids containing viral DNA do not accumulate in strains expressing gp210-GFPmut1. The ratio of DAPI staining in the nucleus compared to the cytosolic regions on either side of the nucleus that typically contain bouquets revealed a significant decrease (*p < 0*.*0001*, n ≥ 200) in bouquet staining relative to nuclear staining when gp210-GFPmut1 was expressed, compared to GFPmut1 alone or mutant gp210(H82R)-GFPmut1 (Figure 2B). In contrast, ΦPA3 infection morphology and bouquet formation is undisturbed by the overexpression of gp210 (Figure S2B).

**Figure 2.**
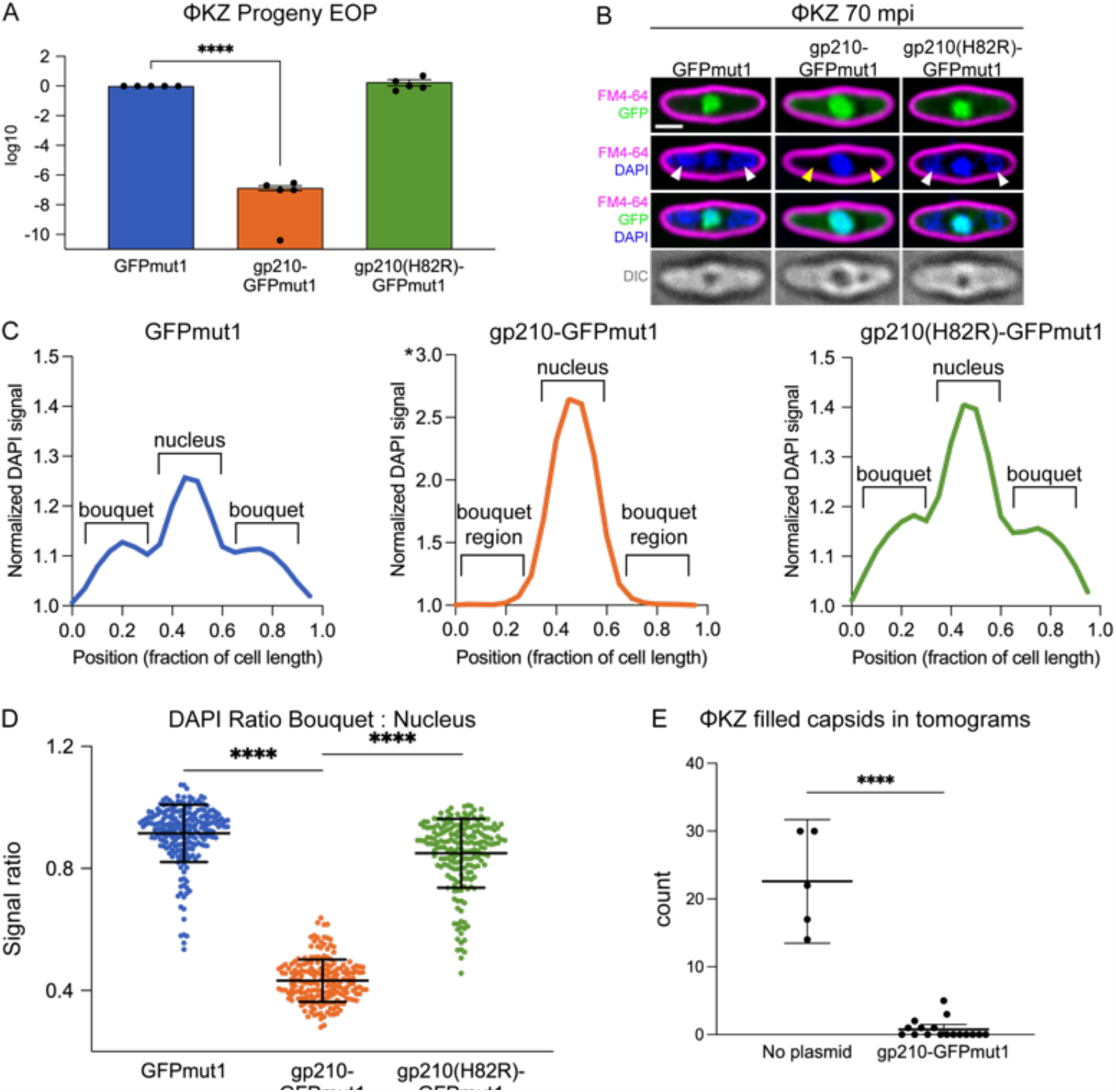
Gp210-GFPmut1 results in a loss of ΦKZ progeny and selects for point mutations of ΦKZ gp178. (A) Live fluorescence microscopy of ΦKZ infections stained with FM4-64 (magenta: membranes) and DAPI (blue: DNA) 70 minutes post infection (mpi). Infections proceeded in the presence of either GFPmut1, gp210-GFPmut1, or gp210(H82R)-GFPmut1 (green: GFP). DAPI stained capsids in phage bouquets labeled by white arrowheads. Absence of DAPI stained capsids in cells expressing gp210-GFPmut1 noted with yellow arrowheads. DIC: Differential Interference Contrast. Scale bar is 1 μm. (B) Ratio of DAPI signal in the bouquet compared to the nucleus at 70 mpi. *Unpaired t-test*, **** *p < 0*.*0001*. Error bars represent standard deviation. GFPmut1 n=220, gp210-GFPmut1 and gp210(H82R)-GFPmut1 n=200. (C) Line plots of DAPI signal intensity along a bisecting line at 70 mpi, average of n=50. *note that the y-axis for gp210-GFPmut1 is double that of the other panels. (D) EOP of ΦKZ progeny collected from washed infections of cells expressing the indicated protein, measured by spot titer on cells without plasmid. Progeny grown in the presence of gp210-GFPmut1 plaque with an efficiency of 0.00001% of the progeny grown with GFPmut1 (*p<0*.*0001*, n=5). Progeny produced with gp210(H82R)-GFPmut1 have a relative EOP of 190% (n=5). Error bars represent SEM and a paired t-test was used. (E) Number of capsids filled with DNA observed in tomograms of the control strain (no plasmid, n=5) or the strain expressing gp210-GFPmut1 (n=17) at 90 mpi.

To test if viable ΦKZ progeny are produced from infections containing gp210-GFPmut1, cells were infected with phage, washed to remove unbound parent phage, and the host cells were allowed to lyse and release the progeny for collection after 2 hours. The ΦKZ progeny lysate produced in cells expressing gp210-GFPmut1 plated with an efficiency of 0.000014% (*paired t-test, p < 0*.*0001, n = 5*) compared to the progeny produced with only GFPmut1 and this was rescued by the H82R mutation of gp210 (Figure 2D). Taken together, these results indicate that gp210-GFPmut1 prevents the accumulation of capsids with DNA and more broadly, the production of infectious virions.

The loss of infective progeny virions (Figure 2D) and the lack of DAPI-stained phage bouquets (Figures 2A-C) led us to hypothesize that gp210 cleavage of the gp178 gene disrupts virion production. To visualize the macromolecular organization of a ΦKZ infection in the presence of gp210-GFPmut1, we performed cryo-focused ion beam milling coupled with cryo-electron tomography (cryo-FIB-ET). The ΦKZ infections in *P. aeruginosa* without any plasmid contained some empty capsids and many complete capsids full of DNA, with an average of 22.6 per tomogram (± 3.3 SEM, n=5, Figure 2E) that were grouped into non-spherical bouquets by 90 mpi (Figures 3A-B and Figure S3C). However, when gp210-GFPmut1 was expressed (Figures C-D), only a few filled capsids were observed (magenta arrows, Figure 3D), with an average of 0.76 per tomogram (± 0.34 SEM, n=17, Figure 2E), demonstrating a 96.6% decrease in virion production (Figure 2E, *unpaired t-test, p < 0*.*0001*) in agreement with our fluorescence microscopy results (Figures 2A-C).

**Figure 3.**
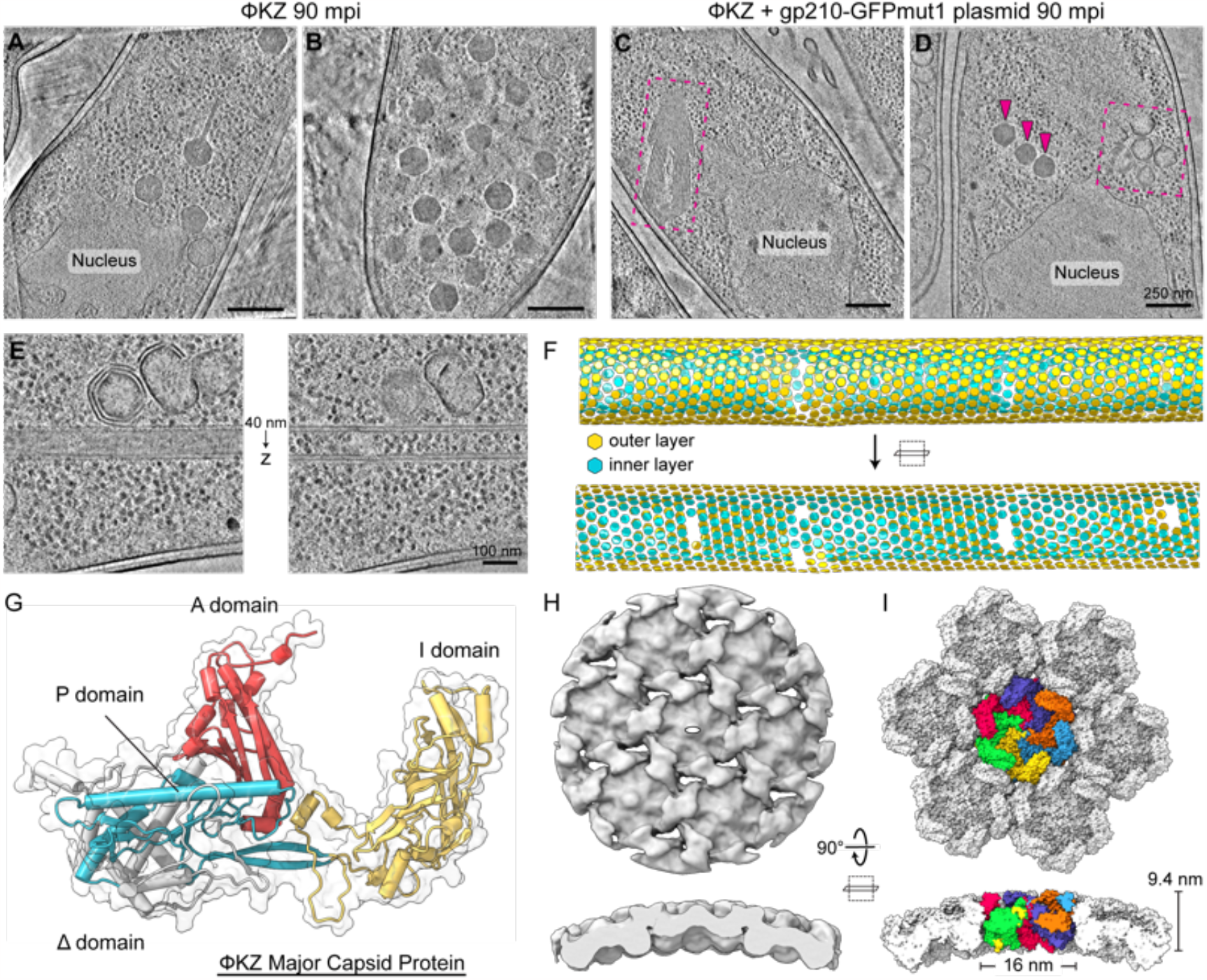
Targeting of ΦPA3 homing endonuclease to ΦKZ phage nucleus results in misassembly of ΦKZ major capsid protein. (A, B) Cryo-FIB-ET of ΦKZ-infected *P. aeruginosa* cells at 90 mpi. (C, D) Cryo-FIB-ET of ΦKZ-infected *P. aeruginosa* cells expressing ΦPA3 gp210-GFPmut1 at 90 mpi. Magenta arrows point to a few properly assembled ΦKZ capsids and magenta dashed boxes indicate regions containing misassembled capsomer complexes. (E) Tomogram slices of a double-layered capsomer tube separated by 40 nm in the Z-direction. (F) Lattice plot of aligned subtomograms from the region depicted in (E). Subtomograms from the outer and inner layers are colored yellow and cyan, respectively. (G) AlphaFold2/ColabFold predicted structure of the ΦKZ major capsid protein (MCP) depicted as a cartoon model with overlaid transparent surface representation. The structure is annotated according to the HK97-fold features. (H) Exterior face and slabbed side views of the subtomogram reconstruction of the ΦKZ MCP from the tubular assemblies with a white oval indicating the 2-fold symmetry. (I) Same views as in (H) of the surface representation of the fitted model of the ΦKZ MCP protomer into the subtomogram reconstruction with the central hexamer protomers colored individually and surrounding hexamers colored white. Scale bars A-D: 250 nm, E: 100 nm

Utilizing cryo-FIB-ET of infected cells expressing gp210-GFPmut1, we observed a significant deficit of filled capsids and also discovered many unusual structures that have never before been observed during nucleus-forming phage infection (magenta dashed boxes, Figures 3C and D). These unusual structures span a range of shapes, sizes, and number of layers. Some assemblies conform to capsid-like geometries, while others are tubular structures of variable diameter (Figures 3E and F). Regardless of shape, the layers of all these structures possess a similar texture and thickness of ∼9.4 nm (Figures 3H and I), suggesting a common elementary unit. Through subtomogram analysis of the tubular structures, we obtained an approximately 13 Å resolution map in which we could readily fit a predicted structure of the immature ΦKZ major capsid protein (MCP, gp120) to explain the density (Figures 3E-I and S4). The immature status of the MCP is marked by the presence of the N-terminal Δ-domain, which is proteolytically cleaved upon proper capsid maturation^32^ (Figures 3G and S5). Previous work on the P22 and T4 MCP describe similar types of geometries (e.g., irregularly sized capsids, spiral lattices, and tubular “polyheads”) when the initial capsid nucleation and scaffolding process is disrupted^33–35^. These results show that gp210 targeting of the ΦKZ genome disrupts capsid assembly.

This work demonstrates that gp210 is a homing endonuclease encoded within an intron interrupting a ΦPA3 vRNAP gene. If gp210 is imported into the ΦKZ nucleus, it is able to cut the vRNAP gene of co-infecting phage ΦKZ at the site homologous to the intron insertion site in ΦPA3 (Figure 4C). This results in the inhibition of ΦKZ virion assembly and prevents subsequent rounds of infection, significantly reducing ΦKZ fitness. Given the observed effects on ΦKZ fitness, we asked whether this mobile intron containing the gp210 homing endonuclease provides an advantage to phage ΦPA3 during competition with ΦKZ. We created an isogenic ΦPA3 lacking gp210 (ΦPA3Δ210) using Cas13a^36^ (Figure S6) and quantified the outcome of ΦPA3/ΦKZ competition after a single round of co-infections. Cells were infected with a ΦPA3 MOI of 10 and a ΦKZ MOI of 0.1, such that most cells infected by ΦKZ would be co-infected with ΦPA3. Cells were washed to remove unbound phage and the number of ΦKZ progeny was determined after one replication cycle. We found that the ΦKZ population size was 3.7-fold lower when competing against wildtype ΦPA3 with gp210 compared to mutant ΦPA3 lacking gp210, while wildtype and mutant ΦPA3 produced equivalent numbers of viable progeny (Figures 4A and B). This 3.7-fold difference in a single generation would amplify over evolutionary time, providing an advantage for ΦPA3. The mobile intron containing gp210 provides a mechanism by which ΦPA3 can defend its niche by decreasing the number of virions produced by the competitor ΦKZ. This work reports the first demonstration of a selective advantage conferred by a mobile intron to the host virus.

**Figure 4.**
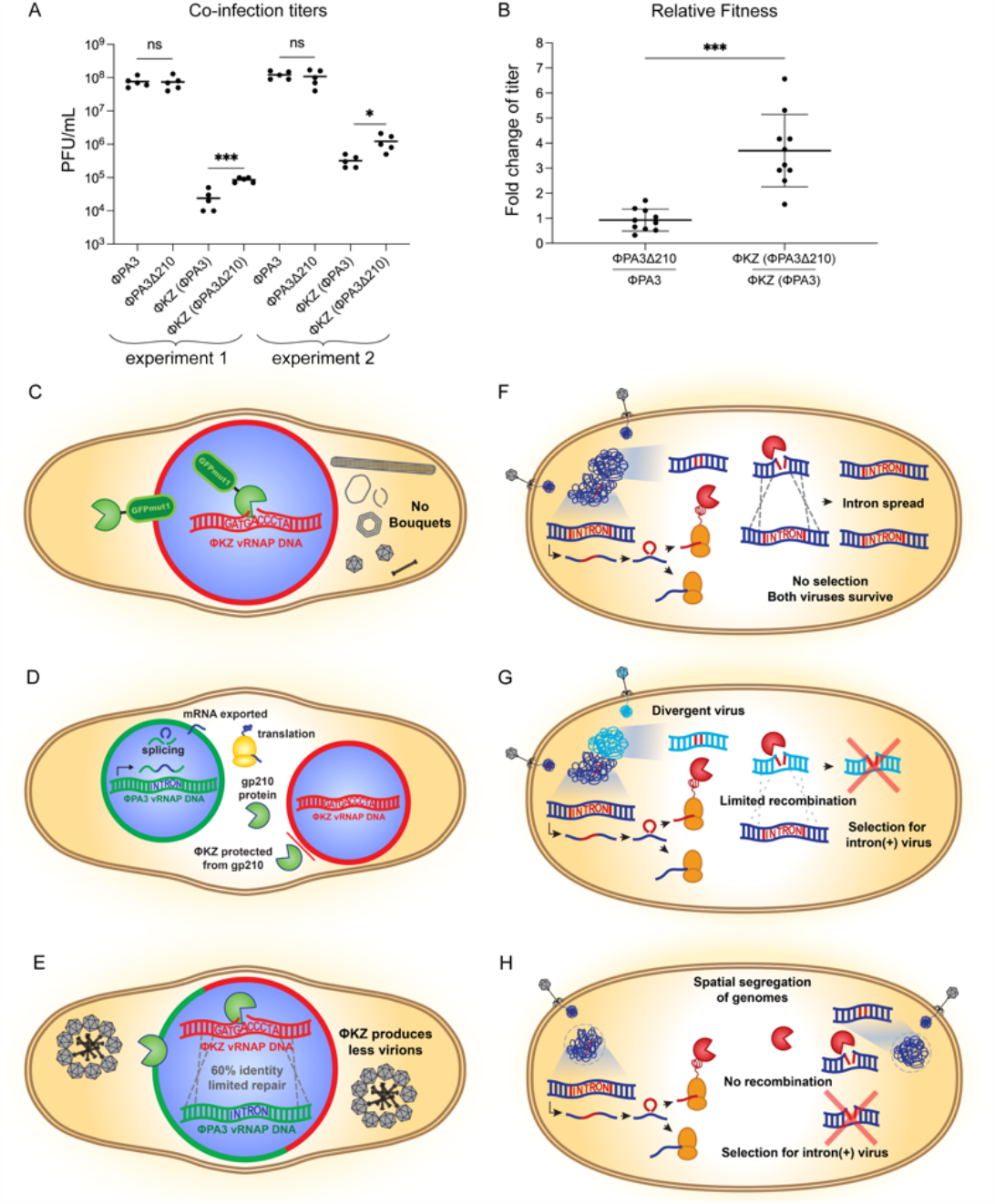
Models of the selective advantage provided by a mobile intron in viral competition. (A) Titer (PFU/ml) of ΦPA3 or ΦKZ after one round of competition with either wildtype ΦPA3 (ΦPA3) or ΦPA3 lacking gp210 (ΦPA3Δ210; Figure S6). Two experiments were conducted with 5 independent replicates each. ΦKZ has significantly higher titers after one round of competition with ΦPA3Δ210 compared to wildtype ΦPA3 (*unpaired t test, * p=0*.*014, ***p=0*.*0003)*. There is no difference in titers when wildtype ΦPA3 competes with ΦPA3Δ210. (B) Fold change of titers from (A) of ΦPA3 (left) or ΦKZ (right) when competing with ΦPA3Δ210 relative to competition with ΦPA3 (*unpaired t test, ***p=0*.*0001)*. Each experimental titer (ΦPA3Δ210, ΦKZ(ΦPA3Δ210)) divided by the mean of the control titers (ΦPA3, ΦKZ(ΦPA3)). (C) gp210 (green pacman) tagged with GFPmut1 is artificially imported into the ΦKZ nucleus (red shell) where it cuts ΦKZ DNA, inhibiting ΦKZ bouquet formation. (D) When separate nuclei are formed by ΦKZ (red) and ΦPA3 (green), ΦPA3 expresses gp210 which is translated in the cytosol and ΦKZ excludes gp210 allowing it to replicate simultaneously with ΦPA3. (E) Hypothesis of the effects of gp210 on a hybrid nucleus containing ΦPA3 and ΦKZ which occurs for ∼25% of co-infections^17^. Gp210 is imported and cuts the ΦKZ gp178 gene. Recombination efficiency is reduced with ∼60% identity between the vRNAP alleles, leading to a competitive advantage for ΦPA3. (F) Model of mobile intron spread during co-infection between any related viruses that replicate their genomes freely in the cytoplasm of the host. (G) Model of mobile intron competition between divergent viruses that can physically mix their genomes but sequence divergence has reduced the efficiency of recombination while a highly conserved site is still targeted by the homing endonuclease, giving a selective advantage to the intron(+) virus. (H) Model of mobile intron competition between closely related viruses that display subcellular genetic isolation. Whether spatially separated or physically isolated by a barrier such as the phage nucleus, the homing endonuclease can target the essential gene in the intron(-) virus but the repair template from the intron (+) virus is sequestered. This results in the inhibition of the intron(-) virus and a strong selective advantage for the intron(+) virus.

## Discussion

Mobile introns are widespread, but how often or under what circumstances do they evolve into interfering agents capable of being deployed during viral competition? Here we identify a viral homing endonuclease that demonstrates the capacity to provide a competitive advantage to the host rather than acting as a purely selfish genetic element. Our results suggest a model in which any conditions that separate two genomes encoding a mobile intron but allow mixing of their encoded proteins, have the potential to involve homing endonuclease-based interference competition. In the case of nucleus-forming phage, the barrier formed by the phage nucleus separates co–infecting phage genomes and thereby segregates the repair template from the targeted genome, allowing for interference competition by the homing endonuclease. The gp210 endonuclease encoded by ΦPA3 is excluded from the ΦKZ nucleus (Figure 4D) but it is expected to be imported into hybrid nuclei formed by both phage (Figure 4E). Within these hybrid nuclei, recombination repair using the ΦPA3 genome is expected to be limited by the amount of divergence between the genomes, explaining the observed competitive advantage. For more closely related nucleus-forming phages that share nuclear import pathways, such as the less-diverged ancestors of ΦPA3 and ΦKZ, acquisition of a homing endonuclease by one phage would confer a strong competitive advantage. During co-infection of these two closely related phages, import of a homing endonuclease into both nuclei results in a cut made in the genome of the phage lacking the intron, which cannot be repaired by the intron-containing genome that is sequestered in the other nucleus. This makes the endonuclease a potent weapon that inhibits closely related co-infecting phages lacking the intron, while also driving the evolutionary divergence of the phage nucleus to exclude these toxic proteins, as is the case for ΦKZ.

Since the general mechanism of homing endonucleases is highly conserved, the principles that lead to a competitive advantage as reported here apply to many other viruses. For example, other mechanisms besides phage nucleus formation that result in spatial separation of genomes occur for viruses in both bacteria and eukaryotic cells^9,53–60^ and are expected to have a similar effect due to a reduced ability to repair the cut made by the nuclease (Figure 4H). For co-infecting viral genomes that replicate in close proximity without a physical barrier, the intron will spread if there is sufficient homology (Figure 4F) but recombination can be limited by sequence divergence^49,50^ (Figure 4G). Mobile introns are known to prefer highly conserved regions of essential genes^61^ so as two genomes diverge, the nuclease target site remains largely unchanged while surrounding DNA loses homology with the competitor, limiting rescue by homologous recombination. In this way, the homing endonuclease of the mobile intron can be deployed against related but divergent phages, even without subcellular genetic isolation. Overall, these models suggest that intron-containing viruses can have a strong fitness advantage during competition and provide a possible explanation for the ubiquity of mobile introns among viruses.

Homing endonucleases with the potential to facilitate interference competition are broadly distributed across diverse phage families, fungi, and archaea^8,9,37–47^. Figure 1B displays a small sampling of endonucleases that are highly conserved with gp210 and found in jumbo phages, which infect a wide variety of host cells including Gram negative and Gram positive bacteria isolated from many environments^48^, suggesting that the general principles outlined here are broadly at work in ecosystems across the planet. In addition, every mobile intron has the potential to evolve as a weapon, due the differences in rates of divergence between conserved target sites and the surrounding genome^49,50^. This mechanism is especially important in the evolution of viruses due to their constant competition through co-infection^51,52^ and rapid rates of viral replication where even small selective advantages are quickly amplified over time. More broadly, this new understanding of the fitness advantage provided by weaponized mobile introns could come into play under the right circumstances for any intracellular genetic competition, including between plasmids^62^, viruses, and hosts.

## Materials and Methods

### Bacterial strains, phage, and growth conditions

*P. aeruginosa* PAO1 derivative K2733 was used as the host in all experiments. It was grown at 30°C in LB and 15 μg/ml gentamycin was used for plasmid selection. Phage stocks of ΦKZ and ΦPA3 were collected from medium titer plate lysates using phage buffer (10 mM Tris (pH 7.5), 10 mM MgSO_4_, 68 mM NaCl, and 1 mM CaCl_2_) and stored at 4°C with titers of 10^12^ and 10^10^, respectively.

### Plasmid construction and transformation

Plasmids were either constructed by Gibson assembly^17^ or synthesized and cloned by Genscript (see Table S2). The vector for all plasmids was pHERD-30T^17–19,65^ and Genscript cloned the inserts using SacI and SalI. Plasmid transformation was accomplished by 2 kV electroporation of K2733 cells washed with 300mM sucrose and stored at −80°C.

### Generation of gp210-Deficient mutant ΦPA3

*P. aeruginosa* K2733 cells containing Cas13a expression vector pHERD30T_LbuCas13a_PhiPA3gp210_HNN_Asn1g1 with guide targeting the HNN endonuclease active site of gp210 (TCTAAGTTATCAAGGCGGTCATCGCCTTTTA) were grown to OD= ∼0.5 in LB containing 15 μg/mL gentamicin and 1% arabinose. 100 μL of culture were infected with ∼10^9^ PFU of ΦPA3 and mixed with 5mL of LB 0.2% arabinose 0.35% agar cooled to 55°C. Lawns of bacteria expressing gp210-targeting Cas13a were grown by plating this mixture on LB 15 μg/mL gentamicin 1.5% agar. Individual plaques containing escape mutants were selected and streak-purified three more times on lawns of bacteria expressing gp210-targeting Cas13a. The region of the genome targeted by Cas13a was then amplified by PCR and the amplicon sequenced by Sanger sequencing to determine mutation.

### ΦPA3 ΦKZ co-infection and quantification

*P. aeruginosa* K2733 was grown in LB and diluted to an OD 0.1. 1 mL of culture was infected with ∼10^9^ PFU of either ΦPA3 or gp210 KO ΦPA3 (MOI: ∼10) and ∼10^7^ PFU of ΦKZ (MOI: ∼0.1). Infected cultures were incubated for 5 min at 30°C rotating and then the cells were pelleted at 8,000rcf for 1.5 min. Pellets were washed with 1mL of LB three times and then infected cells were resuspended in 200uL of LB and incubated while rotating at 30°C for 90 minutes (∼1 infection cycle). Remaining cells were pelleted and supernatant containing phage was removed and serially diluted from 10^−1^ to 10^−8^. Total phage concentration was determined by spot titer on lawns of K2733. ΦKZ concentration was determined by spot titer and whole plate titer on lawns of K2733 expressing Cas13a targeting the start codon of ΦPA3 chimallin (pHERD30T_Lbu_PhiPA3-shellg1, TTTGGGTCCTTGTTGGGTCTGTTGCATTTGC). Replicate titers from each experiment were normalized to the mean of the control condition (Wildtype ΦPA3, ΦKZ coinfection) from each individual experiment and then combined to determine the mean fold-change in titer between the two conditions.

### Protein expression, purification, and characterization

Full length WT gp210 and mutant genes were cloned into UC Berkeley Macrolab vector 1B (Addgene #29653) to generate N-terminal fusions to a TEV protease-cleavable His_6_-tag. Proteins were expressed in *E. coli* strain Rosetta2 pLysS (EMD Millipore) by induction with 0.25 mM IPTG at 20°C for 16 hours.

For protein purification, cells were harvested by centrifugation, suspended in resuspension buffer (20 mM Tris-HCl pH 7.5, 300 mM NaCl, 5 mM imidazole, 5 mM 2-mercaptoethanol and 10% glycerol) and lysed by sonication. Lysates were clarified by centrifugation (16,000 rpm 30 min), then supernatant was loaded onto a 5 mL Ni^2+^ affinity column (HisTrap HP, GE Life Sciences) pre-equilibrated with resuspension buffer. The column was washed with buffer containing 20 mM imidazole and 100 mM NaCl, and eluted with a buffer containing 250 mM imidazole and 100 mM NaCl. The elution was loaded onto an anion-exchange column (Hitrap Q HP, GE Life Sciences) and eluted using a 100-600 mM NaCl gradient. Fractions containing the protein were pooled and mixed with TEV protease (1:20 protease:gp210 by weight), then incubated 48 hours at 4°C for tag cleavage. Cleavage reactions were passed over a Ni^2+^ affinity column, and the flow-through containing cleaved protein was collected and concentrated to 2 mL by ultrafiltration (Amicon Ultra-15, EMD Millipore), then passed over a size exclusion column (HiLoad Superdex 200 PG, GE Life Sciences) in a buffer containing 20 mM Tris-HCl pH 7.5, 300 mM NaCl, and 1 mM dithiothreitol (DTT). Purified proteins were concentrated by ultrafiltration and aliquoted and frozen at −80°C for biochemical assays. All mutant proteins were purified as wild-type.

### Nuclease activity assays

Plasmids were miniprepped from 5 mL of overnight culture of NovaBlue *E. coli* cells. Purified gp210 was mixed with 500 ng plasmid DNA in a buffer containing 10 mM Tris-HCl (pH 7.5), 25 mM NaCl, 10 mM MgCl_2_, and 1 mM DTT (50 mL reaction volume), incubated 1 hour at 37°C, then separated on a 1.0% agarose gel. Gels were stained with ethidium bromide and imaged by UV illumination.

### Efficiency of plating (EOP) by spot titers

To determine the efficiency of phage plaque formation on bacteria expressing different fusion proteins, previously published methods were followed^17^. Protein expression was achieved with the indicated fusions by inducing with 1.0% arabinose. For the EOP of progeny, the indicated expression protein was expressed during the replication of the progeny and the titers were performed on the *P. aeruginosa* host, K2733, without any plasmid present.

### IC50 growth curves

Growth curves to determine the amount of phage required for suppression of the bacterial culture to 50% of maximum growth (IC50) were performed as described previously^17^. 95 μl of bacterial host at OD_600_ of 0.1 was combined with 5 μl of ten-fold serial dilutions of a ΦKZ lysate across a 96-well plate. The plates were shaken for an initial 40 minutes at 30 °C in a microplate reader (Tecan Infinite MPlex and Tecan Sunrise) after which OD_600_ measurements were taken every 10 minutes, with continuous shaking at 30 °C between the timepoints. The OD_600_ values were averaged and plotted as a growth curve. IC50 is annotated at half of the OD_600_ reached by the strain when grown with only phage buffer. Fusion proteins were expressed by 0.1% arabinose.

### Single cell infection assay

Single cell infections of *P. aeruginosa* were visualized using fluorescence microscopy^17–19,65^. Briefly, 1% agarose pads containing 25% LB, 2 μg/ml FM 4-64, and 1 μg/ml DAPI were inoculated with 5 μl of cells (OD_600_ 0.4) and incubated for 3 hours at 30°C in a humidor and then infected with 10 μl of a high titer phage lysate. Imaging was performed with a DeltaVision Elite Deconvolution microscope (Applied Precision, Issaquah, WA, USA). Images were further processed by the aggressive deconvolution algorithm in DeltaVision SoftWoRx 6.5.2 Image Analysis Program, and analyzed using Fiji 1.53c software.

### Lysis time-lapse

Time-lapse to measure lysis was performed as previously described^17^. Infections were established on agarose pads (described above) without any stains. Beginning after 45 minutes post-infection (mpi), images were captured every 10 minutes for 4 hours using UltimateFocus. The cells that lysed during the imaging were counted and reported as a percentage of the total number of starting cells. The percentages were averaged and plotted for both bacterial host strains.

### Progeny collection

ΦKZ progeny were generated in the presence or absence of functional or nonfunctional ΦPA3gp210. Homogeneous liquid cultures of *P. aeruginosa* K2733 were grown in LB for 3 hours at 30°C. The OD_600_ was measured and cultures were back-diluted to an OD_600_ of 0.05 and induced with 1% arabinose for 1 hour to express the protein of interest (GFPmut1, gp210-GFPmut1, gp210-sfGFP, or gp210H82A-GFPmut1). Cells were grown to an OD_600_ of about 0.1 and infected with ΦKZ at an MOI of 30 to ensure high infection rates. During early infection (30 mpi, after cells would be infected but before lysis begins), cells were centrifuged at 3220 rcf for 10 minutes to pellet infected cells and the supernatant was removed, filtered through a 0.45 μm filter to remove any residual bacterial cells, and saved as the “parent” phage sample. Cells were resuspended in fresh LB and washed twice more by centrifugation and resuspension to remove any residual parent phage from the supernatant. When the infections would be entering later stages (80 mpi), centrifugation was halted to prevent premature lysis of infected cells and resuspended cells were concentrated 5-fold in LB to ensure a high titer of progeny phage and incubated for 2-4 hours at 30°C to allow for complete lysis of infected cells. After incubation, the cells were centrifuged for 1 minute at 21130 rcf to pellet the cells and cell debris, and the supernatant containing the progeny phage lysate was collected and filtered through a 0.45 μm filter to remove any residual bacterial cells. Progeny and parent phages were stored at 4°C.

### Cryo-FIB-ET

Cells expressing gp210-GFPmut1 were grown for 3 hours in a humidor at 30°C on 10 agarose pads (1% arabinose, 1% agarose, 25% LB). 10 μl of high titer ΦKZ lysate was added to the pads and incubated for another hour before 25 μl of 25% LB was added at RT and the cells were gently scraped from the pads using the bottom of an eppendorf tube and collected for plunging.

A custom-built manual vitrification device (Max Planck Institute for Biochemistry, Munich) was filled with liquid nitrogen (LN_2_) to maintain a and an ethane/propane gas mixture was condensed to a liquid in a LN_2_-cooled cup in the device. Room humidity was kept around 30% to mitigate water contamination. Holey carbon-coated QUANTIFOIL® R2/1 copper grids were glow discharged (0.2 mbar, 20 mA, 60 sec) 30 minutes prior to plunging using a Pelco easiGlow™ system. 7 μL of concentrated infected cells were put on the carbon side of each grid. Samples were blotted with Whatman filter paper No. 1 to remove excess liquid and plunge-frozen in the liquid ethane/propane to be vitrified.

Grids were mounted into cryo-FIB AutoGrids (TFS) and milled as previously described using an Aquilos (TFS) dual-beam scanning electron microscope under cryogenic conditions^66^. Briefly, areas of interest were rough-milled with an ion-beam current of 0.1-0.50 nA, followed by fine milling and polishing at 30-50 pA. One grid was prepared for each condition to yield two lamellae for the wild-type infection and four lamellae for infected host cells expressing the gp210-GFPmut1 fusion.

Lamellae were imaged under cryogenic conditions using a Titan Krios (TFS) transmission electron microscope operated at 300 kV and equipped with a K2 direct electron detector (Gatan) mounted post a Quantum 968 imaging filter (Gatan). The microscope was operated in EFTEM using a 20 eV slit-width and the detector operated in counting mode. Data collection was performed semi-automatically using SerialEM-v3.8.0^67^. Tilt-series were collected following a dose-symmetric scheme starting at the specimen pre-tilt and targeting a tilt-range of +/- 60° in increments of either 2°, 2.5°, or 3°, using a pixel size of either 3.457 Å or 4.265 Å. Exposure times for each tilt-movie were adjusted to maintain constant counts on the detector throughout the tilt-series. The cumulative dose for each tilt-series was usually between ∼150-200 e/Å^2^ and defocus between 5 to 7 μm. For tilt-series of the gp210-GFPmut1 expressing host cells, 14 were acquired at 3.457 Å and 3 at 4.265 Å. For the tilt-series of wild-type host cells, 2 tilt-series were acquired at both pixel sizes.

All tilt-movies were motion-corrected and defoci estimated using Warp-v1.09^68^. Tilt-series alignment was performed by patch-tracking using Etomo from IMOD-v4.10.28^69^. Tomograms were reconstructed using Warp-v1.09 with the deconvolution filter applied to those used for visualization.

For each condition, ribosome template-matching was performed using Warp-v1.09 using EMD-25183 filtered to 50 Å as a reference and results curated with Cube (https://github.com/dtegunov/cube). Unless otherwise specified, all conditions were aligned and classified similarly using RELION-v3.1.3^70,71^. Initially selected subtomograms were reconstructed at 12 Å per pixel and refined against the template-matching reference. Reference-free 3D-classification without alignment was then performed to remove false-positives, followed by an additional round of refinement. For further refinement of particle alignment and tilt-series parameters, half-maps were upsampled to the original pixel size and imported into M-v1.0.9^72^. Lastly, subtomograms were reconstructed at the original pixel size and subjected to an additional round of reference-free 3D-classification and refinement in RELION-v3.1.3. For the gp210-GFPmut1 expressing host cells, this procedure resulted in a ribosome map at 9.9 Å from 15,439 subtomograms from data collected at 3.457 Å and a map at 13 Å from 4,439 subtomograms from the data collected at 4.265 Å. For wild-type host cells, this procedure resulted in a ribosome map at 39 Å from 921 subtomograms from the data collected at 3.457 Å and a 13.7 Å map from 6,475 subtomograms from data collected at 4.265 Å.

For the tubular assemblies of the ΦKZ major capsid protein, an initial reference was generated using the MATLAB-v2019b (Mathworks) implementation of Dynamo-v1.1514^73^. The surface of an individual, single-layer tube was twice over-sampled assuming a repeat distance of 145 Å (Figure S4A,B). Subtomograms were extracted at 12.5 Å per pixel and oriented normal to the surface with random in-plane rotation. These subtomograms were aligned against a smoothed version of the average generated from the unaligned subtomograms (Figure S4D). The alignment procedure restricted out-of-plane searches to a 60° cone, allowed unrestricted in-plane searches, restricted shifts to 75 Å from the initial position, applied an ad hoc lowpass filter of 40 Å, and C1 symmetry each iteration for 20 iterations. After 15 iterations, the reference ceased to change in subsequent iterations and showed clear hexamer densities with apparent C2 symmetry (Figure S4D,E). The reference was shifted to center on a hexamer, cropped, and C2 symmetrized to use for alignment of the remaining subtomograms.

To more precisely define the subtomogram position from the variable tube diameter and layers, the tube axes were first manually traced and refined. For each tubular structure, the start/end points were manually annotated and subtomograms were defined as overlapping segments along the tube axis. The subtomograms were then aligned to a smoothed reference generated from the unaligned subtomograms. The alignment procedure restricted out-of-plane and in-plane rotations to 15°, restricted shifts to 75 Å from the initial position along the tube axis but only 25 Å perpendicular to the axis, applied an ad hoc lowpass filter of 40 Å, and C1 symmetry each iteration for 5 iterations. A moving average using a sliding window of 3 points along the aligned subtomogram positions was computed as the refined tube axis. Radii were determined from line-profiles of Z-projected, rotationally averaged final averages. The tube axis and radii determined through this procedure were used to extract subtomograms as described for the initial reference. Any subtomogram positions outside of lamellae boundaries were discarded prior to alignment.

All subtomograms were aligned against the initial, symmetrized reference generated above while applying a cylindrical mask to exclude adjacent layers in the case of multi-layer tubes (Figure S4F). This alignment was performed for a single iteration while restricting out-of-plane searches to a 60° cone, restricting in-plane searches to 180°, restricting shifts to 75 Å from the initial position, applying an ad hoc lowpass filter of 40 Å, and C2 symmetry.

Post-alignment subtomograms were subject to a multi-step curation procedure. First, aligned subtomograms which converged to the same position were dereplicated by discarding the one with the lower cross-correlation score. Next, subtomograms with fewer than 5 neighboring subtomograms within 125 to 162.5 Å were discarded. Then, the subtomogram cross-correlation scores were normalized on a per-tomogram basis based on the subtomograms’ orientation with respect to the missing-wedge to account for systematically lower scores of these orientations (Figure S4C), as previously described^74^. The subtomogram positions were evaluated using the PlaceObject-v2.1.0 plugin^75^ for UCSF-Chimera-v1.14^76^ to determine suitable cross-correlation threshold to exclude misaligned subtomograms. Threshold values typically fell between 0.5-0.6 based on the normalized values. This resulted in 21,121 curated subtomograms.

Subtomogram metadata was converted for further processing with Warp/RELION using dynamo2m-v0.2.2^77^. Subtomograms were extracted at 3.457 Å using Warp-v1.0.9 and further refined with C2 symmetry in RELION-v3.1.3 using half-sets composed on a per-tube basis. The final subtomogram average obtained has estimated resolution of 13 Å (Figure S4K). Final aligned subtomogram positions were assessed by a neighbor plot^78^, which reflected the expected geometry based on the obtained subtomogram average (Figure S4J).

A coordinate model for ΦKZ major capsid protein was predicted using the ColabFold^79^ implementation of AlphaFold2^80^. Multiple copies of the resulting model were docked into the final subtomogram average and flexibly fitted into the map using the Namdinator webserver^81^. Geometry minimization with restraints generated from the original AlphaFold2 prediction and C2 symmetry constraints was performed using Phenix-v1.19^82^ (Figure S4G). Rendering of the maps, models, and surface properties was performed using ChimeraX-v1.4^83^.

### Isolation and sequencing of ΦKZ mutants resistant to gp210

To obtain ΦKZ mutants that were able to plaque successfully when the host was expressing gp210-GFPmut1, whole plate infections were generated from 100 μl of cells (OD_600_ ∼0.4) induced for over an hour with 0.1% arabinose and combined with 10 μl of ΦKZ lysate (10^12^). Phage adsorption was allowed to proceed while stationary for 10 minutes at room temperature (RT) prior to the addition of 5 ml warm 0.35% LB top agar. The mixture was poured onto an LB plate with 15 μg/ml gentamycin and incubated overnight at 30 °C in a humidor. Isolated plaques from each plate were streaked onto a new plate and overlaid with top agar containing 0.1% arabinose and cells expressing gp210-GFPmut1 then incubated overnight. Plaque isolation was repeated twice more and the final isolate was streaked to achieve web lysis. Those plates were soaked with 5 ml chilled phage buffer for 5 hours then collected and centrifuged at 3,220 rcf for 10 minutes at 4°C. The supernatant was filtered through a 0.45 μm Corning membrane filter by syringe, 5 drops of chloroform were added, it was shaken by hand for 2 minutes, then centrifuged again to separate the aqueous phase. Lysate was drawn off the top for standard titers on cells expressing GFPmut1 or gp210-GFPmut1. For mutants displaying resistance to gp210, 2 whole plate infections for each isolate were generated and the lysates collected as described above.

Phage genomic DNA was isolated from each mutant using 10 ml lysate incubated with 5 μl of each RNaseA (100 mg/mL) and DNaseI (20mg/mL) at 37°C for 30 minutes. Then 4 ml of phage precipitant solution (30% PEG 8000, 19.3% NaCl in ddH_2_O) was added and incubated overnight at 4°C. Samples were centrifuged at 10,000 rcf 4°C for 20 minutes and the pellets were resuspended in 0.5 ml sterile water for incubation at 5 minutes at RT. Then 2.5 ml of Qiagen Buffer PB was added and incubated at RT for 10 minutes with occasional swirling. The resuspensions were filtered through Qiagen PCR Purification columns and washed twice with Qiagen Buffer PE. The columns were dried with an extra 2-minute spin before 100 μl of 37°C Tris-EDTA buffer was added and allowed to soak for 5 minutes before elution.

Samples were sequenced by the Microbial Genome Sequencing Center (MiGS) in Pittsburgh. Whole genome sequencing was performed using the Illumina NextSeq 2000 platform at a depth of 200Mbp. MiGS provided paired end reads (2x151bp) and reported variations from the Genbank entry for ΦKZ (NC_004629.1).

### Image quantitation

To measure differences in DNA content of the phage nuclei using DAPI concentration, cells expressing either GFPmut1 or gp210-GFPmut1 were grown with 1% arabinose and 1 μg/ml DAPI for 3 hours then infected by ΦKZ for 45 minutes. Raw images were analyzed in Fiji 1.53c. The raw integrated density of each phage nucleus was measured by an inscribed circle on the Z slice closest to the middle of the nucleus. Background raw intensity was measured from empty space next to each infected cell that was measured. Intensity was normalized to the area of the region measured and the background subtracted. A violin plot of these values was generated using Prism 9.2.0 by GraphPad.

## Author Contributions

E.A.B., C.M., T.G.L., R.K.L., A.P., S.R., G.N.M, J.L., E.A., and S.S. conducted experiments. E.A.B., C.M., T.G.L., and K.D.C. analyzed data and created figures. E.A.B. and J.P. conceptualized the original manuscript. J.R.M., K.P., E.V., K.D.C., and J.P. supervised the work, provided feedback on the manuscript, and secured funding.

## Acknowledgements

This work was supported by an Emergent Pathogens Initiative grant from the Howard Hughes Medical institute (to E.V., J.P., K.D.C., and J.R.M.), the National Institutes of Health R01-GM129245 (to J.P., K.P., and E.V.) and R35 GM144121 (to K.D.C.) and the National Science Foundation MRI grant NSF DBI 1920374 (to E.V.). E.V. is an investigator of the Howard Hughes Medical Institute. T.L. was supported by a Simons Foundation Award of the Life Sciences Research Foundation. We acknowledge the use of the UC San Diego cryo-EM facility, which was built and equipped with funds from UC San Diego and an initial gift from the Agouron Institute. We would like to thank Avani Mylvara and Livia Songster for their participation in experiments.

The authors declare no conflict of interest.

## Data Availability

The final subtomogram average of the ΦKZ major capsid protein from this study has been deposited with the Electron Microscopy Data Bank with accession number EMD-40674.

