## Supplemental Figures for "A mobile intron facilitates interference competition between co-infecting viruses"

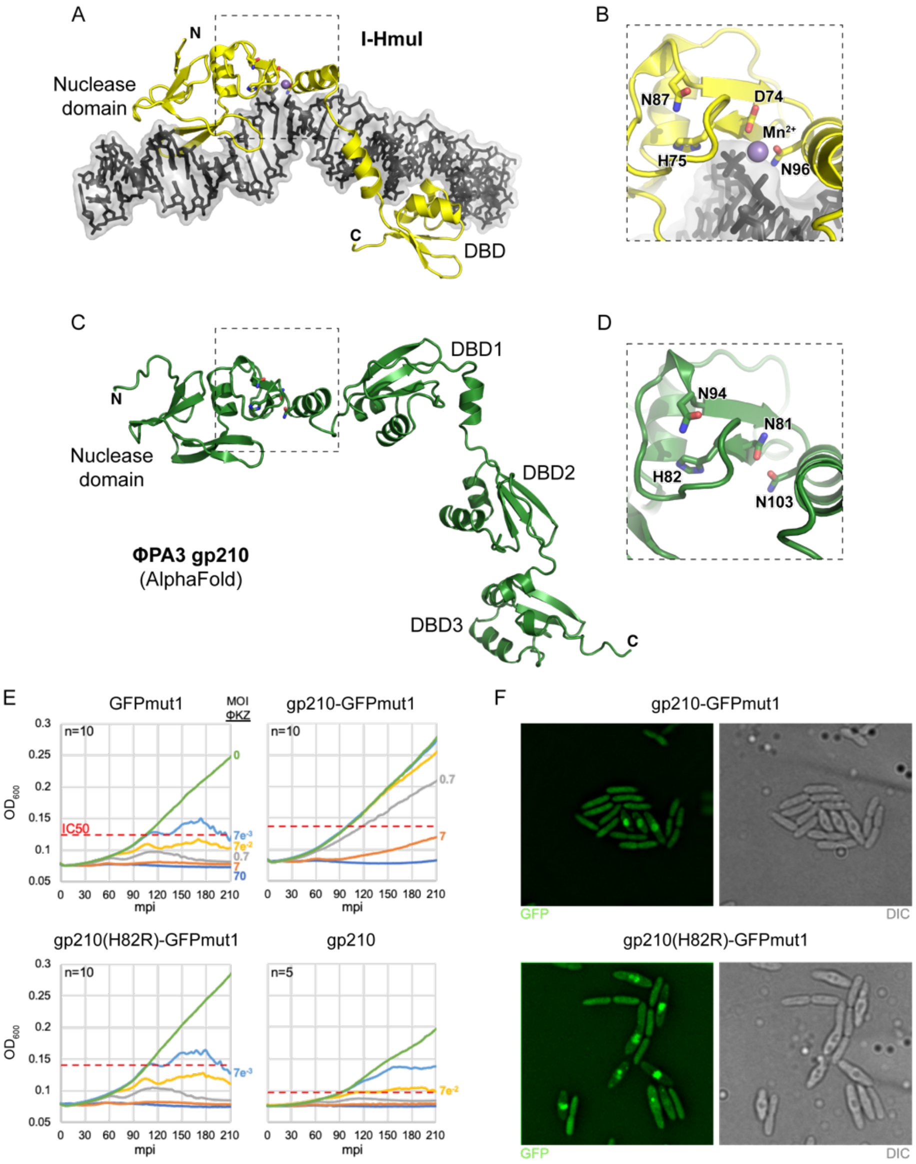


**Figure S1. ΦPA3 gp210 is a putative homing endonuclease.**
(A) Structure of the I-HmuI homing endonuclease from phage SPO1 (yellow) bound to its target DNA site (black). The protein has two domains: an N-terminal HNH/N nuclease domain, and a C-terminal DNA binding domain (DBD). (B) Closeup of the I-HmuI active site, with catalytic residues shown as sticks and a bound $Mn^{2+}$ ion shown as a purple sphere. (C) AlphaFold-predicted structure of ΦPA3 gp210 (green), with predicted N-terminal HNH/N nuclease domain and three C-terminal DNA binding domains (DBD1-3) labeled. (D) View of the ΦPA3 gp210 active site in the same orientation as panel (B), with putative active site residues shown as sticks. (E) ΦKZ growth curves measuring $OD_{600}$ of bacteria in liquid culture showed an MOI of $7e^{-3}$ was required to achieve 50% inhibition of cell growth (IC50: red dotted lines) when cells were expressing GFPmut1 (n=10). gp210-GFPmut1 increased the IC50 MOI to 7, a 1,000-fold decrease in ΦKZ fitness (n=10). gp210(H82R)-GFPmut1 rescued the IC50 back to $7e^{-3}$ (n=10), while untagged gp210 in the cytoplasm caused only a 10-fold increase in required MOI (n=5). (F) Field images of ΦKZ infecting *P. aeruginosa* cells expressing either gp210-GFPmut1 or gp210(H82R)-GFPmut1 showing that both GFP fusions (green) are imported into the ΦKZ nucleus.

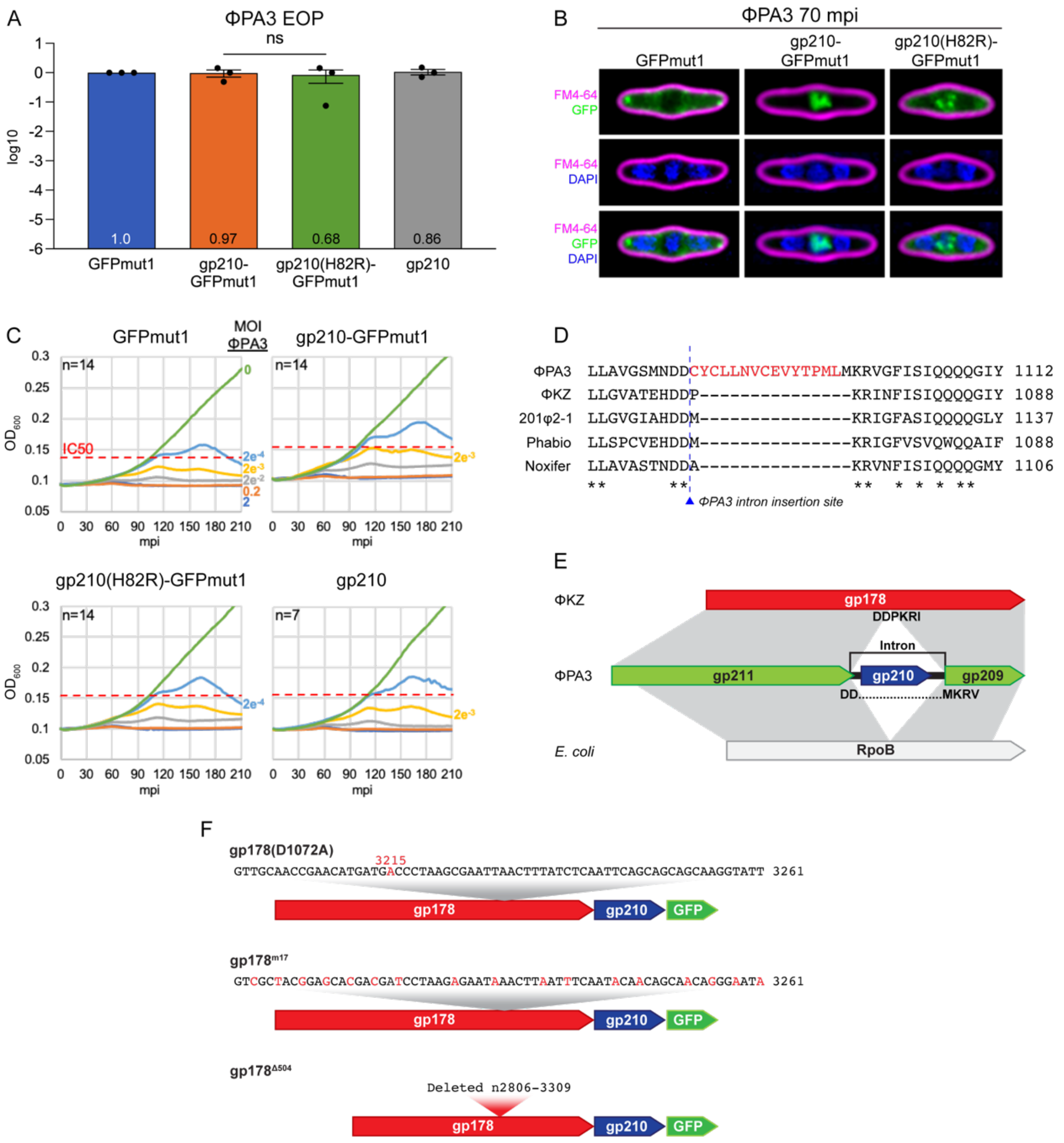


**Figure S2. ΦPA3 is unaffected by gp210. The mobile intron interrupts the ΦPA3 homolog of the ΦKZ gp178 gene.**
(A) ΦPA3 is not significantly affected by expression of gp210-GFPmut1 (*p=0.74*), gp210(H82R)-GFPmut1 (*p=0.51*), or untagged gp210 (*p=0.93*). The difference between gp210-GFPmut1 and gp210(H82R)-GFPmut1 is also insignificant (*p=0.42*). Ratio paired t-tests were used to determine p values. (B) Live fluorescent microscopy of stained ΦPA3 infections in the presence of either GFPmut1, gp210-GFPmut1, or gp210(H82R)-GFPmut1, with no obvious morphological differences between them. (C) ΦPA3 growth curves demonstrating only a 10-fold decrease in ΦPA3 fitness with the expression of gp210-GFPmut1 or untagged gp210 but no change with gp210(H82R)-GFPmut1. (D) Protein alignment of *Pseudomonas* jumbo phage RNAPs reveals an extra 15 residues are included in the Genbank annotation of ΦPA3 gp211. Correct splicing is presented in Figure 2D. (E) Diagram of RNAP subunit genes of nucleus-forming *Pseudomonas* jumbo phages ΦKZ and ΦPA3 aligned with RpoB of *E. coli*. (F) Nucleotide sequences of the gp178 variants: gp178(D1072A) with nucleotide mutation a3215c, gp178$^{m17}$ with 17 silent mutations intended to disrupt gp210 targeting while maintaining the amino acid sequence, and gp178$^{\Delta 504}$ containing an in-frame deletion of 504 bp as a control for the effects of an upstream ORF in the co-expression with gp210-GFPmut1, which does not contain the region targeted by gp210.

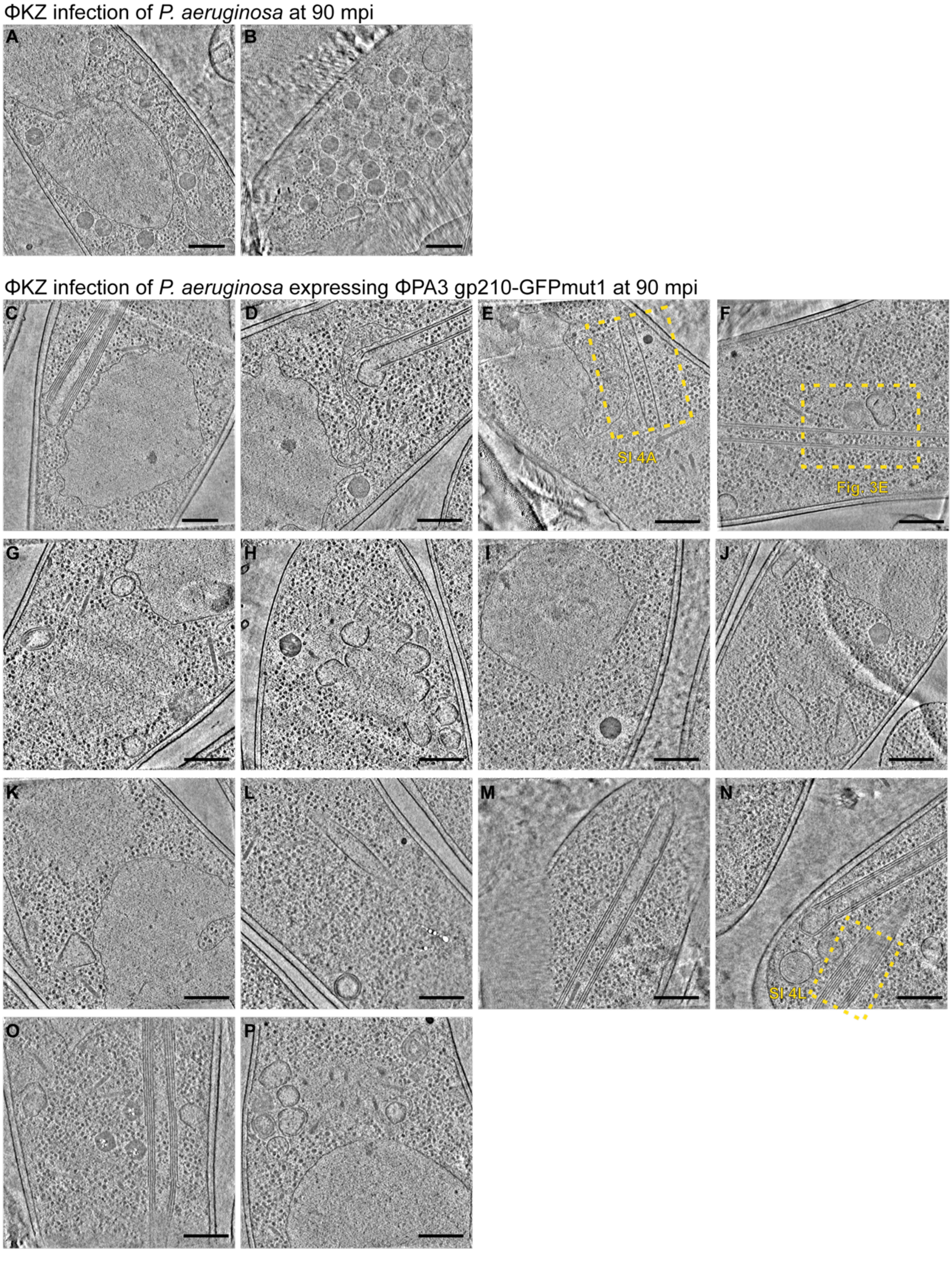


**Figure S3. All collected tomograms of ΦKZ infections.**
(A, B) ΦKZ-infected *P. aeruginosa* cells. (C-P) ΦKZ-infected *P. aeruginosa* cells expressing ΦPA3 gp210-GFPmut1. Regions boxed by yellow dashed lines are enlarged and cropped for display in corresponding figures SI #A, #E, SI #A. Scale bars: 250 nm.

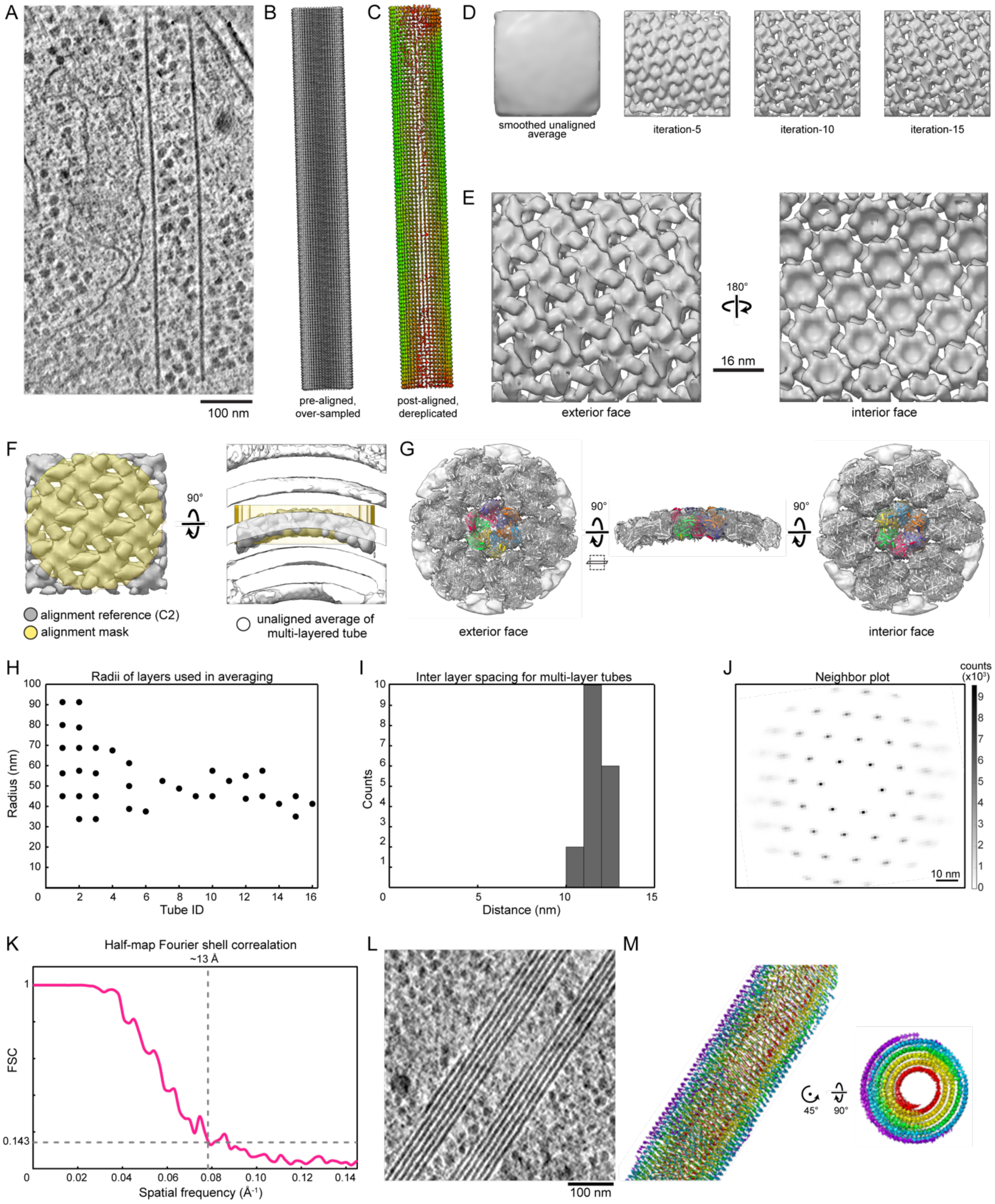


**Figure S4. Subtomogram analysis of tubular assemblies of ΦKZ Major Capsid Protein (MCP).** (A) Exemplar single-layer Major Capsid Protein (MCP) tube used to generate an initial average. (B) Z-arrow lattice plot of over-sampled tube in (A) pre-alignment. (C) Same as (B) post-alignment, dereplicated, and colored red to green for low to high cross-correlation score to the reconstruction, respectively. (D) *Ab initio* reference at indicated iterations of initial alignment. (E) Enlarged views of the exterior and interior faces of the *ab initio* reference. (F) Cropped, shifted reference used to align the remaining data set in grey with alignment mask shown in yellow. Unaligned reconstruction of subtomograms from a multilayer tube is shown white. (G) Views of the final reconstruction with fitted coordinate model of the MCP. The protomers of the central hexamer are colored individually and surrounding hexamers are colored white. (H) Plot of observed tube radii. (I) Histogram of interlayer distances for multilayer tubes. (J) Neighbor plot of center-to-center hexamer distances of the final aligned subtomogram positions. (K) Half-map Fourier shell correlation for the final reconstruction. (L) Tomogram of an aberrant ΦKZ Major Capsid Protein (MCP) assembly observed during expression of ΦPA3 gp210-GFPmut1 in the host cell. (M) Views of the lattice plot depicted as Y-arrows for aligned positions extracted from the region shown in (L). Colors indicate the initially assigned tube layers of the sub tomograms from inner (red) to outer (violet).

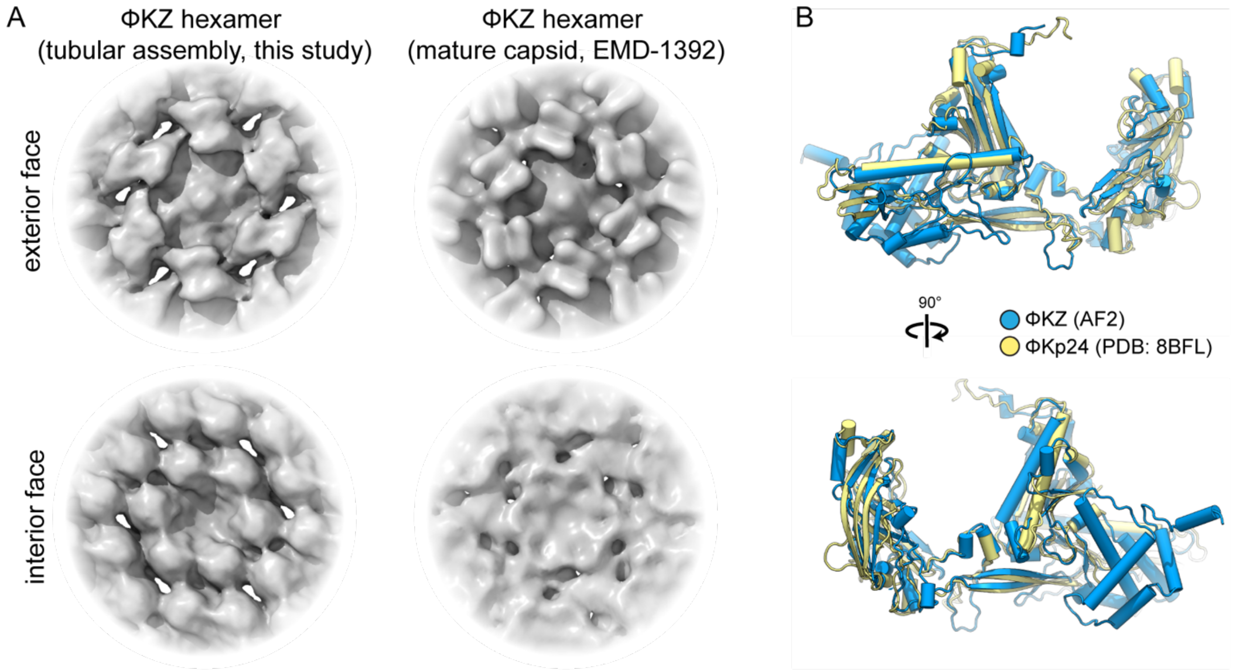


**Figure S5. Comparison of immature ΦKZ and published mature jumbo phage major MCP structures.** (A) Left, exterior and interior faces of the immature ΦKZ MCP density map obtained in this study by subtomogram analysis of the tubular assemblies. Right, exterior and interior faces of the mature ΦKZ density map (EMD-1392) obtained from single-particle analysis of purified virions[63]. (B) Structural alignment of the AlphaFold2 predicted model of the immature ΦKZ MCP (blue) with the experimentally determined structure of the ΦKp24 MCP (yellow, PDB: 8BFL)[64].

A

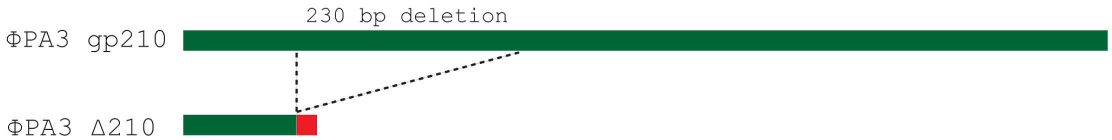


B

ΦPA3 gp210 ATGGCTATAAACTTAAAGGATTTCTCGCCGATTCCTGGTTACAGTAAATATCTTATTTCAAGGGAC...
ΦPA3 Δ210 ATGGCTATAAACTTAAAGGATTTCTCGCCGATTCCTGGTTACAGTAAATATCTTAGA**TAA**CTTAGA...

C

ΦPA3 gp210 MAINLKDFSPIPGYSKYLISRDGD...
ΦPA3 Δ210 MAINLKDFSPIPGYSKYLR*****

**Figure S6. gp210-deficient ΦPA3 isolated with Cas13a guide targeting gp210 catalytic region.** (A) Cas13a escape mutant ΦPA3Δ210 exhibits a 230 base pair deletion leading to a frame shift and early stop codon after amino acid 19. (B) Nucleotide sequence comparison between ΦPA3 gp210 and the ΦPA3Δ210 escape mutant. Identical regions are in green. Mutation region is in red. Early stop codon is bolded. (C) Amino acid sequence comparison between ΦPA3 gp210 and the ΦPA3Δ210 escape mutant. Identical regions are in green. Mutation region is in red. Early stop codon is bolded.
